# HybAVPnet: a novel hybrid network architecture for antiviral peptides identification

**DOI:** 10.1101/2022.06.10.495721

**Authors:** Ruiquan Ge, Yixiao Xia, Minchao Jiang, Gangyong Jia, Xiaoyang Jing, Ye Li, Yunpeng Cai

## Abstract

**Motivation:** The virus poses a great threat to human production and life, thus the research and development of antiviral drugs is urgently needed. Antiviral peptides play an important role in drug design and development. Compared with the time-consuming and laborious wet chemical experiment methods, accurate and rapid identification of antiviral peptides using computational methods is critical. However, it is still challenging to extract effective feature representations from the sequences for the accurate identification of antiviral peptides.

**Results:** This study introduces a novel two-step approach, named HybAVPnet, with a hybrid network architecture to identify antiviral peptides based on neural networks and traditional machine learning methods. Firstly, eighteen kinds of features are extracted to predict labels and probabilities by the neural network classifier and LightGBM classifier, respectively. Secondly, the support vector machine classifier is carried out using the predicted probability of the first step to make the final prediction. The experimental result shows HybAVPnet can achieve better and more robust performance compared with the state-of-the-art methods, especially on independent datasets, which makes it useful for the research and development of antiviral drugs. Meanwhile, it can also be extended to other peptide recognition problems because of its generalization ability.

**Availability and implementation:** The predicted model could be downloaded from: https://github.com/greyspring/HybAVPnet

**Supplementary information:** Supplementary data are available at *Bioinformatics* online.

## 1 Introduction

Viruses have become a great threat to humans and animals because of their high rates of infection and mortality(Calvignac-Spencer, et al., 2021). Viruses can affect all species for long periods of time due to their genetic variation, diversity of transmission, and efficient survival within host cells(Islam and Koirala, 2022). Especially in recent years, the emergence and re-emergence of the current coronavirus disease 2019 (COVID-19) and severe acute respiratory syndrome (SARS) viruses have posed a serious threat to human life and society(Heydari, et al., 2021; Mahmud, et al., 2021). Therefore, it is urgent to develop effective antiviral drugs against various viral pathogens(Saito, et al., 2021). However, traditional treatments often have severe side effects and do not accurately kill viruses. Meanwhile, antiviral drug development is time-consuming and laborious which is not effective enough to address the problem(Hollmann, et al., 2021).

In recent years, drug development based on peptides has attracted wide attention in the industry due to its highly selective, relatively safe, well tolerated and low production costs(Yan, et al., 2022). Antiviral peptides (AVPs), with 8 to 40 amino acids typically(Schaduangrat, et al., 2019), are a promising resource for the treatment of viral diseases. Antiviral peptides can prevent the virus from attaching to or invading the host cell or interfering with viral replication and are easy to synthesis(Basith, et al., 2020). Nowadays, there are some collected, experimentally validated AVP databases(Qureshi, et al., 2015), such as AVPdb(Qureshi, et al., 2014), HIPdb(Qureshi, et al., 2013), APD3(Wang, et al., 2016), CAMP(Thomas, et al., 2010) etc. AVPdb is a comprehensive resource of peptides that have been experimentally validated for antiviral activities. HIPdb is a specific database of experimentally validated HIV inhibiting peptides. Parts of AVPs are collected in the antimicrobial peptide database APD3 and CAMP.

In the past years, many computational tools have been developed to predict AVPs using machine learning methods. AVPpred is the first AVP prediction tool developed using support vector machine (SVM) based on physiochemical properties(Thakur, et al., 2012). Chang KY et al. employed four peptide features and used random forest (RF) classifier to identify AVPs(Chang and Yang, 2013). Zare1 M et al. employed pseudo-amino acid composition (PseAAC) and adaboost with J48 as base classifier to identify antiviral peptides(Zare, et al., 2015). AntiVPP 1.0 selected RF as the final classifier with the new two features relative frequency (Rfre) of all 20 natural amino acids and residues composition of peptides (PEP) to assess the antiviral peptides candidates(Beltran Lissabet, et al., 2019). PEPred-Suite employed an adaptive feature representation strategy to achieve better and robust performance using a two-step feature optimization strategy and eight RF models for eight types of functional peptides, respectively(Wei, et al., 2019). FIRM-AVP achieved a higher accuracy than other models using the informative filtered features from the physicochemical and structural properties of their amino acid sequences(Chowdhury, et al., 2020). Charoenkwan P et al. also comprehensively summarized the above identified tools of AVPs from the feature encoding, classifiers, cross-validation and performance(Charoenkwan, et al., 2021). In addition, deep neural network methods also were employed to extract the high dimensional features for the identification of AVPs from the primary sequence(Li, et al., 2020).

Although the existing methods achieved good performance (Pang, et al., 2021; Timmons and Hewage, 2021), they are not satisfactory for drug development. There are a lot of factors that may improve the model performance, such as unbiased training samples, effective features, model architecture and interpretability, etc.(Agarwal and Gabrani, 2021) In this work, we proposed a novel hybrid network architecture for antiviral peptides identification, named HybAVPnet. To learn the effective features, HybAVPnet is consisted of a two-layer prediction models which are mixed of traditional machine learning models and deep learning models. In the first layer, two neural network and one group of LightGBM classifiers were employed to extract the different aspects of features using one-hot coding, composition, autocorrelation, and profile for amino acid sequences(Yan, et al., 2021). For the second layer, all the probability and label outputs of the first layer were fed into SVM classifier to obtain the final prediction(Vukovic, et al., 2022). The experimental results showed that HybAVPnet could achieve competitive advantages compared with the existing methods.

## 2 Material and Methods

### Datasets

In order to compare our model with other models, we use two groups of datasets from AVPpred. One dataset contains 604 AVPs with experimentally validated antiviral activities and 452 non-AVPs proved to be invalid, which is divided into training and testing subsets, named training set T^544P + 407N^ (544 positive and 407 negative samples) and testing set V^60P + 45N^ (60 positive and 45 negative samples). The another dataset consists of 604 effective AVPs and 604 non-experimental negative peptides from AntiBP2(Lata, et al., 2010), which is also divided into training and testing subsets, named training set T^544P + 544N^ (544 positive and 544 negative samples) and testing set V^60P + 60N^ (60 positive and 60 negative samples). The sequences of AVPs and non-AVPs were statistically analyzed and the amino acid frequency distribution of the datasets was shown in Figure1. It clearly showed that the frequency of amino acid “W” in the positive samples was high. However, there are no obvious rules for the distribution of other amino acids.

**Fig. 1.**
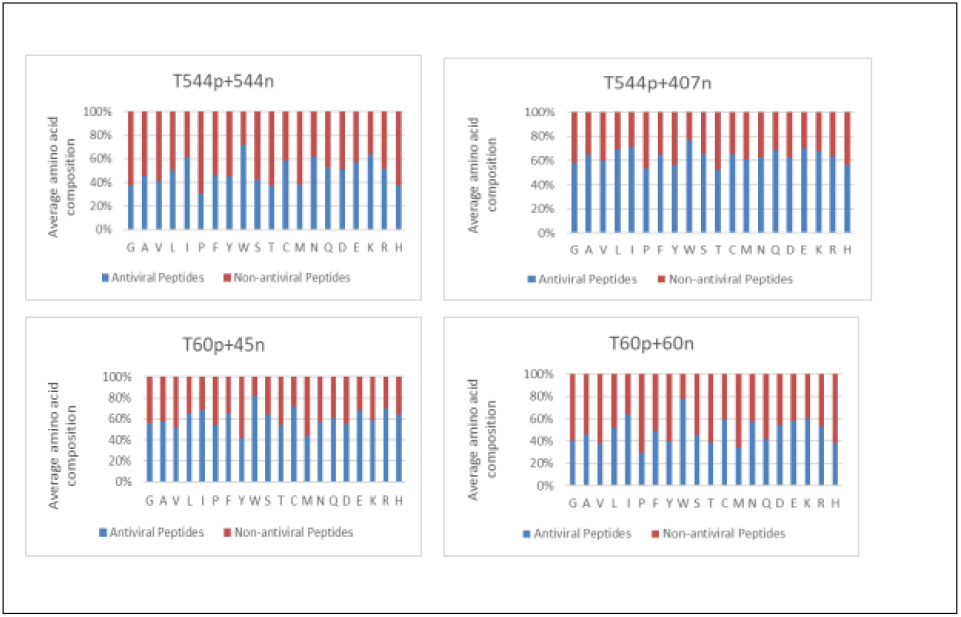
Amino acid frequency distribution of AVPs and non-AVPs. The blue and red bars represent the amino acid frequency distribution of antiviral peptides and non-antiviral peptides respectively.

### Feature Representation

Considering the composition, frequency, physical and chemical properties of the sequence and other information, many features were extracted from the amino acid sequence (Liu, 2019). Among of them, three kinds of features were extracted based on amino acid composition: Basic Kmer (kmer), Distance-based Residue (DR) and Distance Pair (DP)(Liu, et al., 2017). Four kinds of features were generated according to autocorrelation: auto covariance (feature-AC), auto-cross covariance (ACC), cross covariance (feature-CC), and physicochemical distance transformation (PDT)(Jing, et al., 2019). Based on pseudo amino acid composition (PseAAC) and frequency profile, we extracted four and seven kinds of features respectively(Muthu Krishnan, 2018). In total, there are 18 kinds of features which were listed in Table 1. Furthermore, all the features were also input into the neural network to explore the potential relationships between them. In addition, one-hot encoding method in natural language processing was employed to extract the high dimensional features into the neural network structure(Okada, et al., 2019).

**Table 1.** 18 kinds of feature representation methods based on protein primary sequences

| Category | Feature |
| --- | --- |
| Amino acid composition | Basic Kmer (kmer)<br>Distance-based Residue(DR)<br>Distance Pair(DP) |
| Autocorrelation | Auto covariance(feature-AC)<br>Auto-cross covariance(ACC)<br>Cross covariance(feature-CC)<br>Physicochemical distance transformation(PDT) |
| Pseudo amino acid composition | Parallel correlation pseudo amino acid composition(PC-PseAAC)<br>Series correlation pseudo amino acid composition(SC-PseAAC)<br>General parallel correlation pseudo amino acid composition(PC-PseAAC-General)<br>General series correlation pseudo amino acid composition(SC-PseAAC-General) |
| Profile-based features | Select and combine the n most frequent amino acids according to their Frequencies(Top-n-gram)<br>Profile-based Physicochemical distance transformation(PDT-Profile)<br>Distance-based Top-n-gram(DT)<br>Profile-based Auto covariance(AC-PSSM)<br>Profile-based Cross covariance(CC-PSSM)<br>Profile-based Distance-based Top-n-gram(PSSM-DT)<br>Profile-based Auto-cross covariance(ACC-PSSM) |

### Machine Learning Approaches

HybAVPnet identifies antiviral peptides by integrating several machine learning methods, i.e. Light Gradient Boosting Machine (LightGBM), SVM, Convolutional Neural Networks (CNN), and Bidirectional Long Short Term Memory (Bi-LSTM) (Dai, et al., 2021).

In the first layer of HybAVPnet, LightGBM is chosen as the predictor, which is a gradient boosting framework. The LightGBM is based on decision tree algorithms and supports efficient parallel training, with the advantages of faster training speed, lower memory consumption, better accuracy, distributed support, and rapid processing of massive data. SVM is a binary classifier, widely used in the supervised machine learning tasks. It is trying to find the best separated hyperplane in the feature spaces, and maximizes the interval between positive and negative samples on the training set, which makes it different from the perceptron. SVM performs effective in high dimensional spaces. And its kernel can be specified to solve the different problems. CNN (CNN1D) is a kind of feed forward neural network with convolution calculation. It is one of the representative algorithms of deep learning. CNN1D is widely used in sequence models. LSTM is a form of Recurrent Neural Network (RNN), which can take into account the relationship between front and back. So it is often used in sequence model. Bi-LSTM is a combination of forward LSTM and backward LSTM.

### Computational Model

The framework of the whole model HybAVPnet is shown in Figure2, which is composed of three sub models: Neural Network1, LightGBM and Neural Network2.

**Fig. 2.**
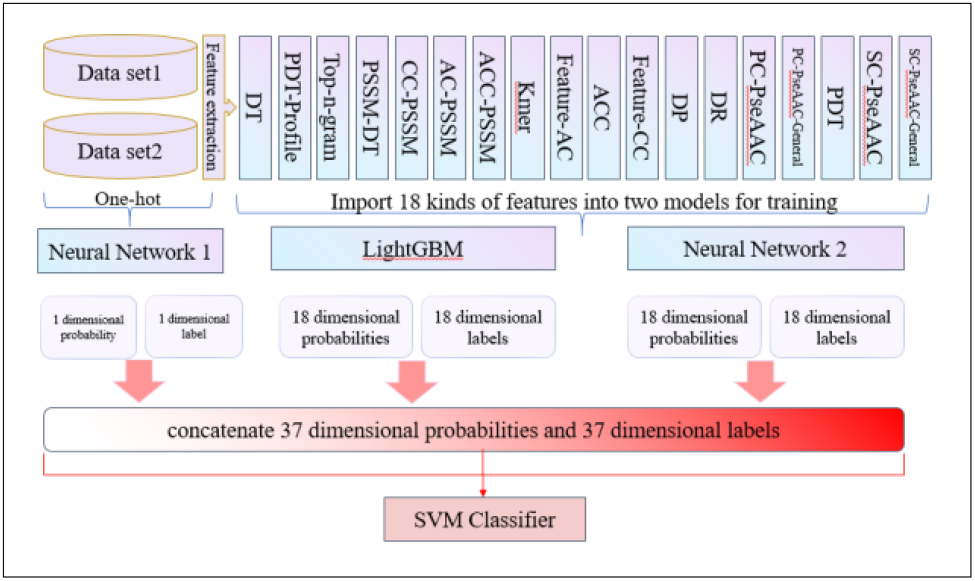
The HybAVPnet model architecture.

Considering that amino acid sequence has its related characteristics, we adopt a series of feature extraction methods to obtain a total of 18 kinds of features. Each kind of features are trained and predicted through LightGBM predictor and Neural Network2 to obtain the initial predicted results. Meanwhile, the amino acid sequence is vectorized according to the specific one-hot coding form, and trained and predicted by the Neural Network1 (Figure 3).

**Fig. 3.**
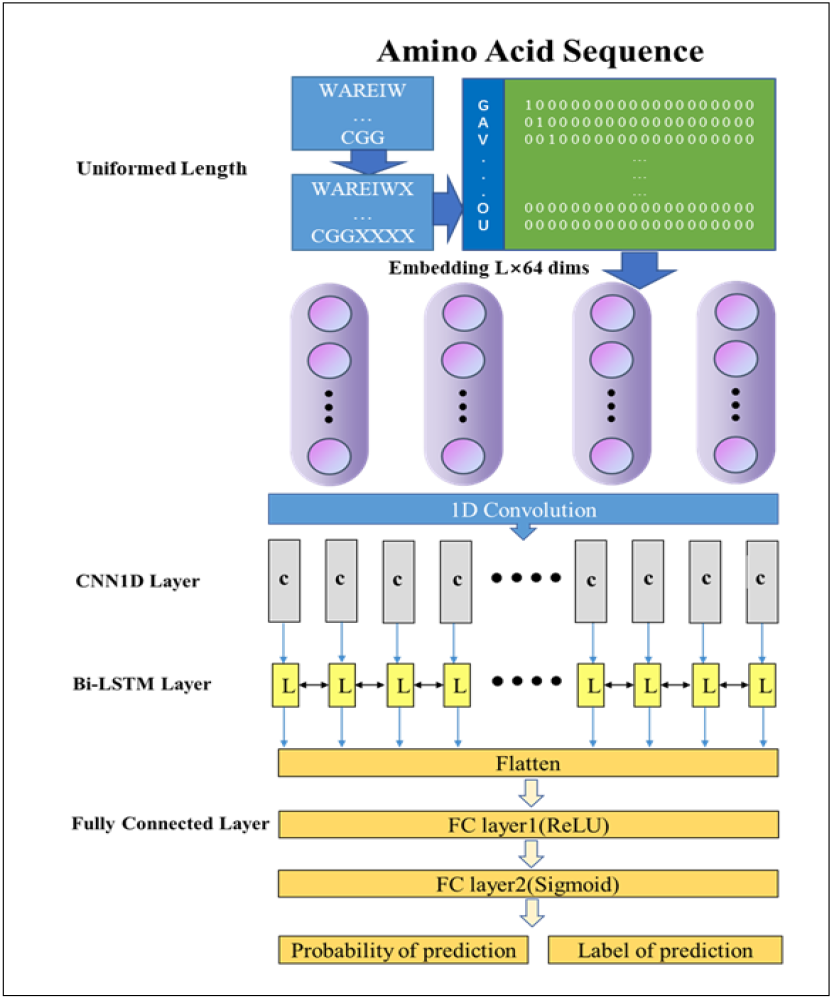
The Neural Network 1 structure. Firstly, each protein sequence is unified the same lengths and encoded by one-hot coding. Then the coded features are imported into neural network as a training set. Through the embedding layer, each vector is converted into 64 dimensions (embedding (input_ dimensions = 26, output_ dimensions = 64, input_length = 1000)). Then the vectors are inputted into Conv1d (filters = 32, kernel_size = 1, activation =‘relu’, strings = 1)). Finally, the outputs of Conv1d are imported into Bi-LSTM layer (bidirectional (LSTM (64, return_sequences = true)). Then, the network obtains the predicted labels and probabilities through two fully connected layers. For the Neural Network 2, we directly import the data into convolution layer and omit the embedding layer.

HybAVPnet consists of two parts, of which the first part includes three sub-models. The first sub-model unifies the protein sequences with different lengths into a certain length. Then, the sequences are vectorized according to the specific one-hot coding form. The coded vectors are inputted into Neural Network1 (Figure 3) to obtain its classification probabilities and classification labels. The second sub model inputs the extracted 18 kinds of features into the LightGBM classifier for training and classification, and achieves the 18 dimensional classification probabilities and classification labels. The third sub model also inputs 18 kinds of features into Neural Network 2 similar to Neural Network1 (omitting the Embedding layer) for training and classification, and gets the 18 dimensional classification probabilities and classification labels. Finally, the obtained 74 dimensional vector datasets are concatenated and put into SVM classifier as training set for the final classification.

The network architecture of Neural Network 2 is similar to the Neural Network 1. Neural Network 2 omits the embedding layer and inputs 18 features directly into the convolution layer. Through the above steps, we obtained a series of initial prediction results. Considering that the factors of predicted probability may have a great impact on the final results, both the probabilities and labels are inputted into the next layer of the network architecture. Therefore, a total of 74 dimensional data from the classification probability and classification labels of three sub models is used as the training set for the next classifier.

In the last layer, some machine learning classifiers are evaluated to find the optimal solution, here we focuses on SVM, LightGBM, Bayes, Decision tree, KNN. Through the comparative experiments, SVM is selected as the final classifier.

### Performance Evaluation

In the experiments, the following metrics were employed to verify the prediction performance of HybAVPnet, including Receiver Operating Characteristic curve (ROC), Sensitivity (Sn), Specificity (Sp), Accuracy (Acc), and the Matthews correlation coefficient (MCC)(Mei, et al., 2020). Five-fold cross-validation and independent test were conducted to evaluate the model on different datasets.

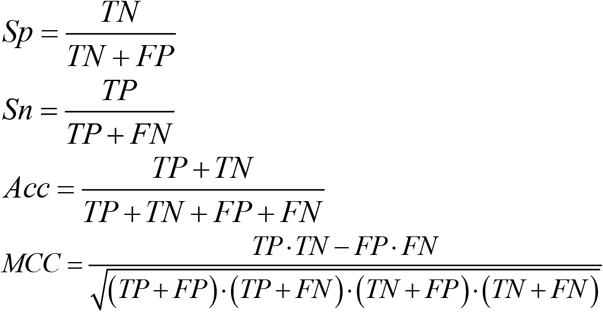

Where TP, FP, TN and FN indicate the number of true positives, false positives, true negatives, and false negatives respectively.

## 3 Results and Discussion

### 3.1. Comparison with the Existing Methods

Five-fold cross-validation was involved to evaluate the model in the training datasets T^544P + 407N^, T^544P + 544N^ and the testing datasets V^60P + 45N^, V^60P + 60N^. The experimental results show that HybAVPnet performs significantly better than other models in T^544P + 407N^, T^544P + 544N^, V^60P + 45N^ and V^60P + 60N^ datasets. In the dataset V^60P + 60N^, HybAVPnet is slightly lower than DeepEvo by 1.7% on sensitivity. DeepPhy and DeepEvo are two different dual-channel deep neural network ensemble models of DeepAvp method.To sum up, HybAVPnet achieves the best performance compared with other existing models in terms of evaluation on crossvalidation and testing datasets as shown in Table 2. Compared with other direct classification models, the classification method combining initial prediction maybe obtain better performance, such as DeepAvp and HybAVPnet. On the datasets T^544P + 407N^ and V^60P + 45N^, the performances of most predicting models are not good except for our method HybAVPnet.

**Table 2.** Comparison of HybAVPnet with existing methods on five-fold cross-validation and independent test datasets. The bold fonts indicate the best results.

| Data set | Model | Acc | Sn | Sp | MCC |
| --- | --- | --- | --- | --- | --- |
| T<sup>544P+407N</sup> | AVPpred | 85.0 | 82.2 | 88.2 | 0.70 |
|  | Chang et al's method | 85.1 | 86.6 | 83.0 | 0.70 |
|  | AntiVPP 1.0 | - | - | - | - |
|  | DeepPhy | 88.0 | 85.5 | 79.7 | 0.65 |
|  | DeepEvo | 83.5 | 84.6 | 82.1 | 0.66 |
|  | **HybAVPnet** | **93.08** | **90.82** | **96.2** | **0.86** |
| T<sup>544P+544N</sup> | AVPpred | 90.0 | 89.7 | 90.3 | 0.80 |
|  | Chang et al's method | 91.5 | 89.0 | 94.1 | 0.83 |
|  | AntiVPP 1.0 | - | - | - | - |
|  | DeepPhy | 88.5 | 88.0 | 89.0 | 0.77 |
|  | DeepEvo | 90.1 | 89.3 | 90.8 | 0.80 |
|  | **HybAVPnet** | **95.83** | **94.17** | **97.34** | **0.92** |
| V<sup>60P+45N</sup> | AVPpred | 85.7 | 88.3 | 82.2 | 0.71 |
|  | Chang et al's method | 89.5 | 91.7 | 86.7 | 0.79 |
|  | AntiVPP 1.0 | - | - | - | - |
|  | DeepPhy | 80 | 83.3 | 75.6 | 0.59 |
|  | DeepEvo | 87.60 | 90.00 | 84.40 | 0.75 |
|  | **HybAVPnet** | **93.27** | **95.00** | **90.91** | **0.86** |
| V<sup>60P+60N</sup> | AVPpred | 92.5 | 93.3 | 91.7 | 0.85 |
|  | Chang et al's method | 93.0 | 91.7 | 95.0 | 0.87 |
|  | AntiVPP 1.0 | 93 | 87 | 97 | 0.87 |
|  | DeepPhy | 89.2 | 88.3 | 90 | 0.78 |
|  | DeepEvo | 93.30 | **96.70** | 90.00 | 0.87 |
|  | **HybAVPnet** | **96.61** | 95.00 | **98.28** | **0.93** |

### 3.2. Ablation Experiments

In the selection of sub-model combinations, ablation experiments were conducted to determine the best combination. The SVM classifier is adopted as the last layer for each model.

Four different models are analyzed with LightGBM, Neural Netwok 1, Neural Network 2, and the fused model HybAVPnet. The average values of the evaluated indicators for the five experiments are taken as the final experimental results. The final results of the ablation experiments are shown in Table 3. It can be found that the results of the fused model HybAVPnet are better than other models whether on the testing set V^60P+45N^ or V^60P + 60N^ after parameter optimization of SVM.

**Table 3.** Comparison of three sub models and joint model in two independent test datasets. The bold fonts indicate the best results.

| Dataset | Model | Acc | Sn | Sp | MCC |
| --- | --- | --- | --- | --- | --- |
| $V^{60P+45N}$ | LightGBM | 89.42 | 91.67 | 86.36 | 0.78 |
|  | Neural Network 2 | 88.27 | 94.67 | 83.48 | 0.78 |
|  | Neural Network 1 | 75.58 | 86.00 | 61.36 | 0.49 |
|  | HybAVPnet | **93.27** | **95.00** | **90.91** | **0.86** |
| $V^{60P+60N}$ | LightGBM | 94.92 | **95.00** | 94.83 | 0.90 |
|  | Neural Network 2 | 94.75 | 93.33 | 96.21 | 0.90 |
|  | Neural Network 1 | 85.59 | 85.67 | 85.52 | 0.71 |
|  | HybAVPnet | **96.61** | **95.00** | **98.28** | **0.93** |

It can be seen from Table 3 that in the testing set V^60P + 45N^, HybAVPnet performs better than other models in all evaluated indicators. However, in the testing set V^60P + 60N^, compared with the LightGBM model, HybAVPnet leads it by 1.69% in accuracy, 3.45% in terms of specificity, 0.03% in terms of MCC, and the same in terms of sensitivity. Therefore, the combined output results of the three sub-models are chosen as the final experimental results for the input of the next SVM classifier. The final experiments prove that the predicted probability has an important impact on the final classified evaluation. So we choose to integrate the predicted probabilities and the predicted labels into the final model. The results prove the fused model may have a strong sense of discrimination in the identification of antiviral peptides.

### 3.3. Comparison with the Different Classifiers

After the pre-classification of the three sub-models in the first step, 74 dimensional initial predicted results were used as the new training set. In the selection of the classifier in the second step as shown in Table 4, a few of traditional machine learning classifiers were adopted to analyze their performance using SVM, Random Forest, LightGBM, Bayes, Decision tree, and KNN classifiers on V^60P+45N^ and V^60P + 60N^ datasets.

**Table 4.** Comparison of the different classifiers. The bold fonts indicate the best results.

| Data set | Model | Acc | Sn | Sp | MCC |
| --- | --- | --- | --- | --- | --- |
| $V^{60P+45N}$ | SVM | **93.27** | 95.00 | **90.91** | **0.86** |
|  | Random Forest | 85.58 | 93.33 | 75.00 | 0.70 |
|  | LightGBM | 81.73 | 86.67 | 75.00 | 0.62 |
|  | Naive Bayes | 80.96 | **97.66** | 58.19 | 0.63 |
|  | Decision Tree | 81.15 | 87.00 | 73.18 | 0.61 |
|  | KNN | 90.77 | 93.00 | 87.73 | 0.81 |
| $V^{60P+60N}$ | SVM | **96.61** | **95.00** | **98.28** | **0.93** |
|  | Random Forest | 95.25 | 94.67 | 95.86 | 0.91 |
|  | LightGBM | 87.29 | 78.33 | 96.55 | 0.76 |
|  | Naive Bayes | 96.27 | **95.00** | 97.59 | **0.93** |
|  | Decision Tree | 87.97 | 89.00 | 86.89 | 0.76 |
|  | KNN | 96.10 | **95.00** | 97.93 | **0.93** |

From Table 4, we can see that on the testing set V^60p+ 45n^, the performance of SVM and KNN is much better than other classifiers. Compared with KNN, SVM achieves 2.5%, 2%, 3.18% and 0.05 higher respectively in Acc, Sn, Sp and MCC. On the testing set V^60p+ 60n^, SVM performs better than other relatively good classifiers Bayes and KNN by 0.34% and 0.51% in Acc respectively, and similarly well in Sn and MCC. While in Sp, SVM achieves better performances than Bayes and KNN by 0.69% and 0.35% respectively.

Furthermore, Receiver operating characteristic (ROC) curve and Precision-Recall (PR) curve are drawn to evaluate the performance of each methods for intuitive comparison, as shown in Figure 4. AUC represents the area under the ROC curve, which is plotted the true positive rate against false positive rate. AUPR stands for the area under PR curve that is plotted precision against recall. On the independent datasets V^60P+45N^ and V^60P + 60N^, SVM can obtain the best balance in performances compared with Random Forest, LightGBM, Bayes, Decision tree and KNN classifiers, and is selected as the last layer classifier..

**Fig. 4.**
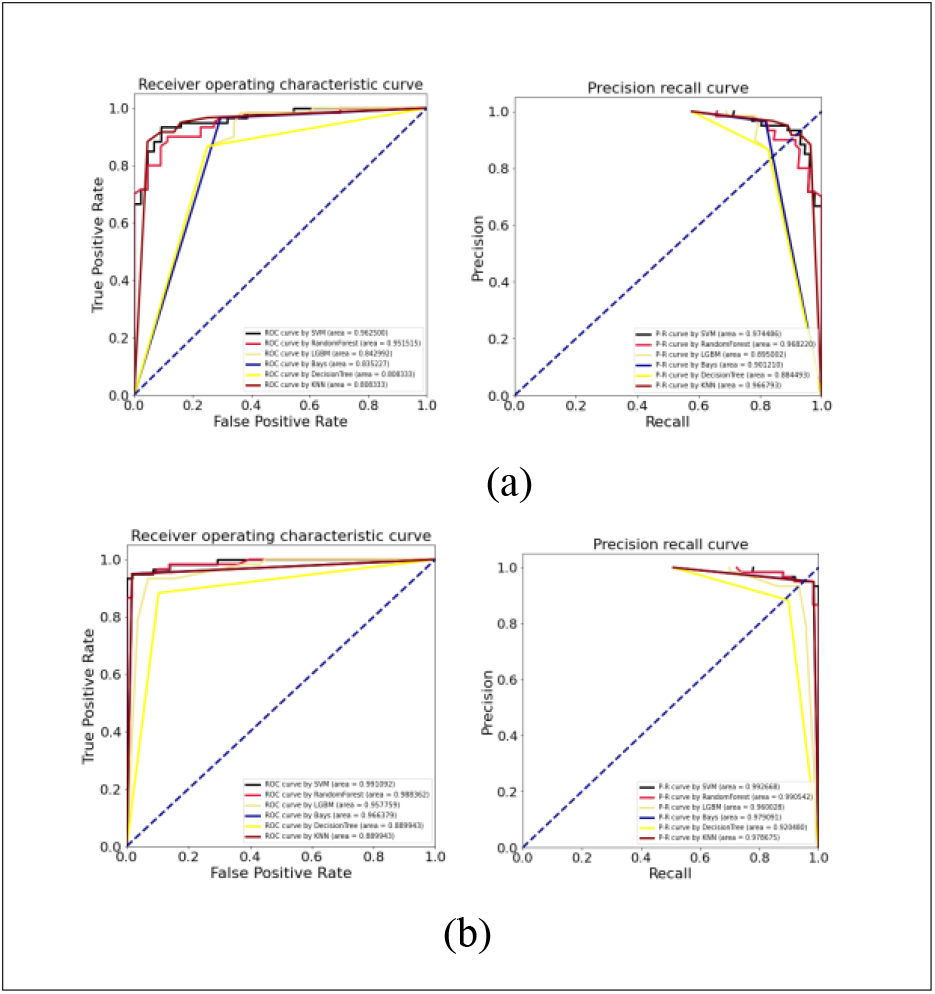
Receiver Operating Characteristic (ROC) and Precision Recall (PR) curve of (a) V^60P+45N^ and (b) V^60P+60N^ datasets.

### 3.4. Visual Analysis

To better interpret the feature representation between the sub models, we adopted t-distributed stochastic neighborhood embedding (t-SNE) to visualize and compare the feature space distribution on V^60P+45N^ and V^60P+60N^ datasets(Kobak and Linderman, 2021).

In the experiment of t-SNE, the LightGBM, Neural Network 1, Neural Network 2 and HybAVPnet are compared to demonstrate the distribution of the new features in the two-dimensional feature space. As shown in Figure 5, the new feature distribution in HybAVPnet is the most efficient and effective compared with other three models to discriminate AVPs from non-AVPs.

**Fig. 5.**
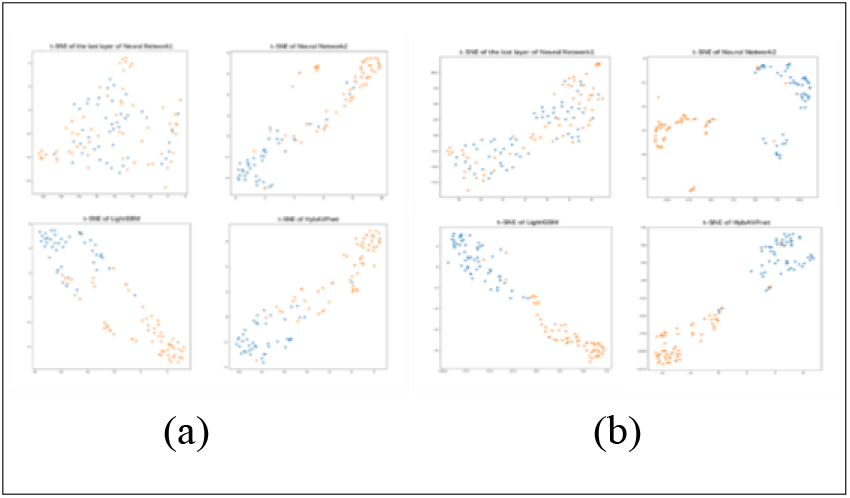
The t-distributed stochastic neighborhood embedding (t-SNE) visualization of (a) V^60P + 45N^ and (b) V^60P + 60N^ datasets. The blue dot represents the distribution of non-antiviral peptides, and the orange dot represents the distribution of antiviral peptides.

## 4 Conclusion

Due to their advantages and good performance, antiviral peptides have potential wide applications in the development of antiviral drugs. To this end, some computational models have been developed to quickly and accurately identify AVPs. In this work, we present a novel hybrid network tool named HybAVPnet to identify AVPs. HybAVPnet takes full advantage of traditional machine learning models and deep learning models to obtain the effective feature representation of amino acid sequences at sequential, structural, and evolutionary levels. Experimental results demonstrated our proposed HybAVPnet model could achieve more discriminative power for the prediction of AVPs and could be easier to separate the positive samples and negative samples. Furthermore, a serial of comparative experiments showed the consistently stability and robustness of HybAVPnet from the five-fold cross-validation and independent test. We except that HybAVPnet can help the development of antiviral peptide drugs and the treatment of related diseases for researches. In the future, we will strive to develop predictive models for various therapeutic peptides to better serve precision medicine.

## Funding

This work was supported by the Zhejiang Provincial Natural Science Foundation of China (No.LY21F020017, 2022C03043), the National key research and development program of China (No.2019YFC0118404), Joint Funds of the Zhejiang Provincial Natural Science Foundation of China (U20A20386), National Natural Science Foundation of China (No. 61702146).

### Conflict of Interest

none declared.

## Notes

### Competing Interest Statement

The authors have declared no competing interest.

